# Single-cell transcriptomics of heterogeneous patient-derived organoids reveals novel therapeutic targets in high-grade serous ovarian cancer

**DOI:** 10.64898/2026.06.05.730209

**Authors:** Bisiayo E. Fashemi, Yukihide Ota, Vijayalaxmi Gupta, Maia L. Elizagaray, Roger Pique-Regi, Nardhy Gomez-Lopez, Mary M Mullen, Dineo Khabele

**Affiliations:** Division of Gynecologic Oncology, Department of Obstetrics and Gynecology, Washington University School of Medicine, and Alvin J. Siteman Cancer Center. St Louis, MO, USA; Center for Reproductive Health Sciences, Department of Obstetrics and Gynecology, Washington University, St Louis, MO, USA; Center for Molecular Medicine and Genetics, Wayne State University School of Medicine, Detroit, MI, USA; Department of Pathology and Immunology, Washington University, St Louis, MO, USA

**Author notes:** **Corresponding Author:** Dineo Khabele, M.D. The Mitchell and Elaine Yanow Professor and Chair of the Department of Obstetrics and Gynecology, Department of Obstetrics & Gynecology. Washington University in St. Louis School of Medicine. St. Louis, MO, 63110, USA.

## Abstract

High-grade serous ovarian carcinoma (HGSOC) is characterized by widespread peritoneal dissemination and poor long-term survival, largely driven by metastatic relapse following initial response to chemotherapy. Defining the molecular programs that enable tumor progression from the primary ovarian site to metastatic niches remains a key challenge. Here, we leverage patient-derived organoids (PDOs) coupled with single-cell RNA sequencing (scRNA-seq) to interrogate tumor evolution and identify regulators of metastatic competence in HGSOC. We profiled PDOs and matched formalin-fixed paraffin-embedded (FFPE) tumor samples from ovarian and omental disease sites across seven patients. Single-cell transcriptomic analysis revealed conserved and patient-specific cellular states and enabled reconstruction of inferred trajectories of tumor progression. Comparative trajectory analysis identified gene expression programs associated with metastatic transition from ovarian to omental tumors. Among these, the heparan sulfate proteoglycan *AGRIN* emerged as a consistently upregulated gene along the metastatic axis. Cell-cell communication analyses suggested that *AGRIN*-mediated signaling involves both epithelial tumor cells and stromal components, implicating the extracellular matrix in shaping metastatic behavior through mechanotransduction and integrin-associated pathways. Functional validation using genetic depletion of AGRIN in ovarian cancer cell lines demonstrated reduced migratory and invasive capacity, supporting a causal role for AGRIN in promoting metastatic phenotypes. Together, these findings identify *AGRIN* as a regulator of metastatic competence in HGSOC and highlight extracellular matrix–associated signaling as a key driver of disease progression. More broadly, this study demonstrates that PDO-based single-cell transcriptomic approaches can uncover actionable regulators of metastasis and provide a scalable framework for therapeutic target discovery across cancer types.

**Significance:** Patient-derived organoids analyzed by single-cell transcriptomics reveal dynamic tumor evolution and uncover *AGRIN* as a regulator of metastatic competence in HGSOC, demonstrating the utility of living tumor models for therapeutic target discovery.

## Introduction

Ovarian cancer is the sixth leading cause of cancer-related deaths for women in the United States [1]. Approximately 75% of patients have high-grade serous ovarian carcinoma (HGSOC), characterized by widespread peritoneal dissemination at diagnosis, and have a five-year survival rate of only 49.1% [2, 3]. Despite initial response to platinum-based chemotherapy, most patients experience relapse driven by metastasis of therapy-resistant tumor cells. Thus, to improve clinical outcomes, we must define the mechanisms by which HGSOC tumor cells interact with the surrounding tumor microenvironment (TME), including stromal, immune, and endothelial compartments, during metastasis.

The HGSOC TME is composed of heterogeneous epithelial, stromal, endothelial, and immune cell populations that collectively shape tumor progression, metastatic dissemination, and therapeutic response. Cancer-associated fibroblasts promote extracellular matrix remodeling and invasion, endothelial cells support angiogenesis and metastatic colonization, and immune populations can adopt immunosuppressive phenotypes that facilitate tumor progression and resistance to therapy. In parallel, epithelial tumor cells exhibit dynamic transcriptional and phenotypic plasticity that enables adaptation to metastatic niches and therapeutic stress. Platinum resistance in HGSOC has also been linked to epithelial plasticity, altered cell-state transitions, enhanced DNA damage repair, and TME-mediated survival signaling, highlighting the need to better define cellular interactions and trajectories within chemotherapy-exposed tumors. Emerging evidence further suggests that reciprocal signaling between tumor and microenvironmental compartments drives metastatic evolution, emphasizing the importance of studying these interactions at single-cell resolution.

An excellent tool for modeling tumor evolution and metastatic behavior *ex vivo* is patient-derived organoids (PDOs). Unlike traditional cell lines, PDOs retain the genetic, cellular, and phenotypic heterogeneity of their parental tumors. Ovarian cancer PDOs have primarily been used to assess therapeutic drug sensitivity [4-8]. However, PDOs can recapitulate key molecular features of cancer metastasis, including epithelial-to-mesenchymal transition (EMT), altered mechanotransduction, TME-derived signaling cues [9-11], and tumor cell plasticity [12-17].

Here, we used HGSOC PDOs to identify new genes involved in metastasis. To do so, we adapted strategies commonly used in used in stem cell and developmental biology studies of non-cancerous organoids. First, we performed single-cell RNA sequencing (scRNA-seq) on PDOs and matched formalin-fixed paraffin-embedded (FFPE) tumor samples obtained from the ovary or omentum of seven patients. Next, we performed transcriptomic and cell trajectory analyses to identify genes whose expression increased along the inferred metastatic pathway from the ovary to the omentum. One such gene was the heparan sulfate proteoglycan *AGRIN*, which is enriched in the extracellular matrix and plays a central role in mechanotransduction, integrin signaling, and regulation of cell migration and invasion in other cancer types. In addition to these structural roles, *AGRIN* has been implicated in regulating immune cell activation and invasion in tumor settings [18-26], suggesting that it may function as a multicellular signaling regulator within the metastatic niche. Our cell-cell communication analysis suggested that *AGRIN*-dependent signaling from both epithelial and stromal compartments contributes to metastatic progression. Finally, genetic depletion of AGRIN in a human ovarian cancer cell line led to decreased migratory and invasive behaviors. Collectively, these findings suggest that *AGRIN* could be targeted to reduce HGSOC metastatic competence and may participate more broadly in extracellular matrix–immune–tumor signaling interactions that shape metastatic progression. Additionally, our strategy could be used to identify genes involved in metastasis in other cancer types for which organoids are available.

## Results

### Transcriptomic analysis of HGSOC patient-derived PDOs and matched tumor FFPEs captures cellular diversity of the TME

We generated patient-derived organoids (PDOs, Figure 1A, Supplementary Figure 1) from four primary ovary and three metastatic omentum tumor samples from patients with stage III or IV high-grade serous ovarian cancer (HGSOC). We decided to focus on better understanding how the TME changed following exposure to chemotherapy. Because of this, all patients had received at least one line of chemotherapy before tumor samples were obtained. We also obtained time-matched formalin fixed paraffin-embedded curls (Figure 1A, Supplementary Figure 1) from the same tumor samples from which the PDOs were derived. Demographic data for the patients are presented in Supplemental Data 1.1.

**Figure 1:**
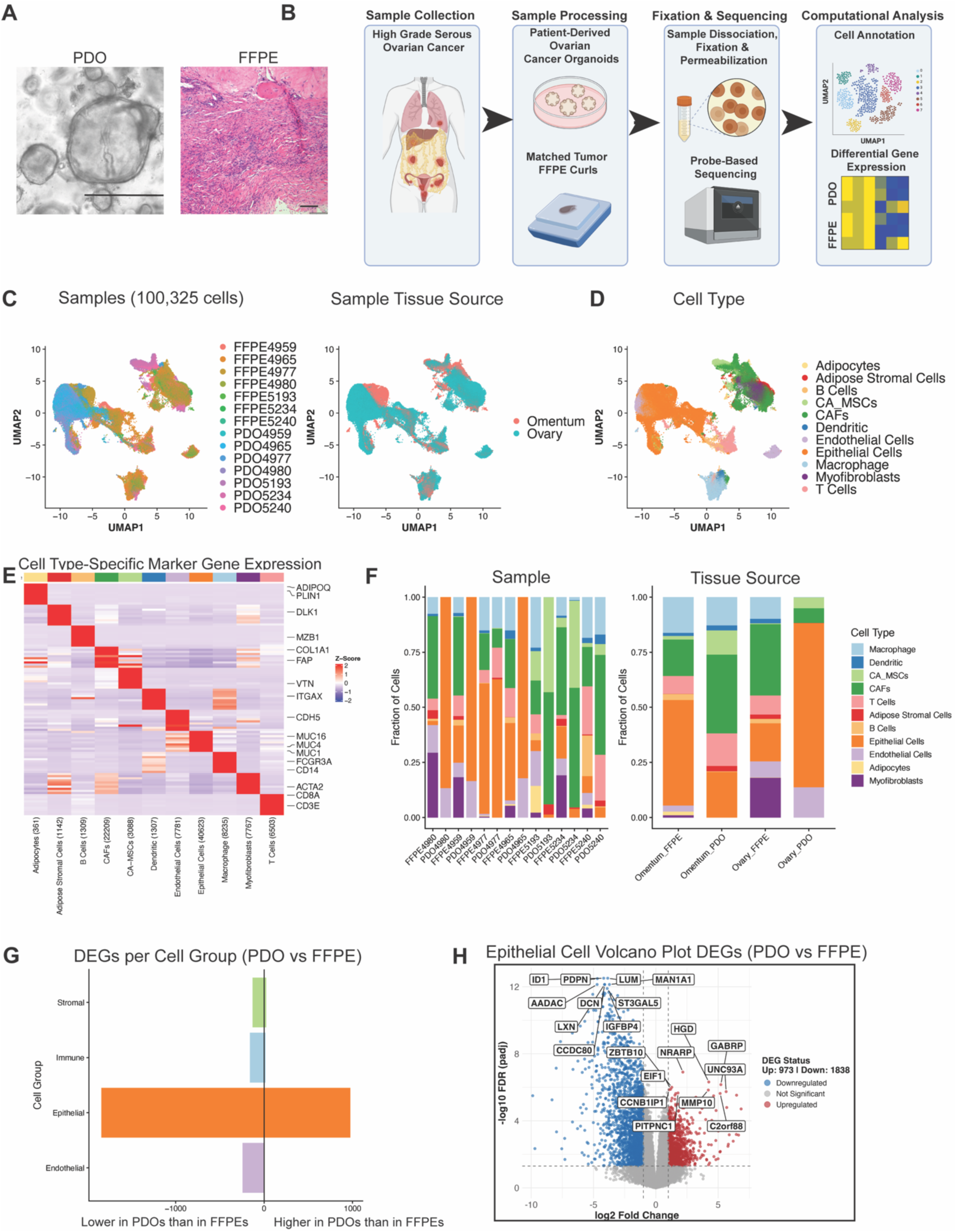
Single-cell sequencing identifies multiple cell types in high-grade serous ovarian cancer patient-derived organoids and matched frozen fixed paraffin-embedded tumor samples. A) Representative brightfield image of a patient-derived organoid (PDO) and hematoxylin and eosin staining image of a matched frozen fixed paraffin-embedded (FFPE) tumor. PDO scale bar = 400 µm, FFPE scale bar = 100 µm. B) Experimental workflow for data collection and analysis. (C-D) Integrated uniform manifold approximation and projection (UMAP) of PDO and FFPE samples (total of 100,325 cells) colored by (C) Samples, Tissue Source, and (D) Cell Type. E) Heatmap showing the top differentially expressed genes for each cell type. Epithelial, stromal, and immune populations were defined according to statistically significant expression of canonical marker genes. F) Stacked bar plots showing the proportion of each cell type per sample and by tissue source. G) Bar plot depicting pseudobulk differential expression analysis comparing stromal, immune, epithelial, and endothelial cells between PDOs and FFPEs. H) Volcano plot highlighting differentially expressed genes in PDO vs. FFPE epithelial cells.

At passage five or earlier, we fixed the PDOs and performed single-cell RNA sequencing (scRNA-seq) on the matched PDO and FFPE samples (Figure 1B). We obtained scRNA-seq data from a total of 58,708 FFPE cells and 41,617 PDO cells (Supplementary Figure 2A-B) and used Harmony to integrate all of the data (Figure 1C). Expression of canonical cell markers revealed 11 major cell types: adipose stromal cells (*SCD*, *PPARG*, *DLK1*), adipocytes (*ADIPOQ*, *PLIN1*), B-cells (*CD79A*, *MS4A1*, *MZB1*), cancer-associated mesenchymal stem cells (CA_MSCs, *ALDH1A3*, *VTN*), cancer-associated fibroblasts (CAFs, *COL1A1*, *ACTA2*, *DCN*, *FAP*), dendritic cells (*CD40*, *ITGAX*), endothelial cells (*VWF*, *CLDN5*, *CDH5*), myofibroblasts (*COL1A1*, *ACTA2*), epithelial cells (*EPCAM*, *MUC1*, *KRT18*), macrophages (*CD14*, *APOE*), and T-cells (*CD3D*, *CD3E*, *PTPRC*) (Figure 1D). In addition to confirming expected tumor-associated stromal and epithelial compartments, the presence of macrophages, dendritic cells, and lymphocytes highlights the contribution of immune populations that are known to influence metastatic niche formation through cytokine production, antigen presentation, and extracellular matrix remodeling. To confirm cell annotation, we identified differentially expressed genes (DEGs) and performed gene ontology (GO) enrichment analysis of the top hits (Figure 1E, Supplementary Data 1.2). T-cells were enriched for T cell receptor signaling pathway term (GO:0050852) and B cells were enriched for B Cell Activation terms (GO:0042113) (Supplemental Data 1.3).

We examined the cellular compositions of individual samples and compared matched FFPE and PDO data (Figure 1F). Overall, FFPE samples exhibited greater cellular diversity than their PDO counterparts. For instance, the ovary-derived FFPE4980 sample contained immune, stromal, epithelial, and endothelial cells, whereas the corresponding PDO4980 sample contained only epithelial and endothelial cells. PDOs derived from omental tumors displayed higher cellular heterogeneity, including immune cells, than PDOs originating from ovarian tumors. The reduced representation of immune populations in PDOs likely reflects selective pressures of *ex vivo* growth conditions and suggests that downstream trajectory analyses primarily capture tumor-intrinsic epithelial state transitions rather than the full complexity of tumor ecosystem evolution occurring *in vivo*.

We next pseudobulked the cells into four groups (stromal, immune, epithelial, and endothelial) and then identified DEGs between PDOs and FFPEs. Epithelial cells exhibited substantially more DEGs than the other cell groups (Figure 1G; Supplementary Figure 3; Supplementary Data 1.4). The top upregulated DEGs in epithelial cells from PDO vs. FFPE samples were associated with proliferation, invasion, and metastasis, and the top downregulated genes were associated with the TME-stroma interaction (Figure 1H, Supplementary Data 1.5). Taken together, these results indicate that the epithelial cell compartment is expanded in PDOs and differs transcriptionally from the epithelial compartment in matched FFPE samples.

### PDO epithelial cell subsets exhibit distinct functional states

Given the pronounced heterogeneity of HGSOC tumors and the large epithelial cell compartment in the PDOs, we wanted to define the functional diversity within this population. We performed consensus non-negative matrix factorization (cNMF) to identify gene expression programs or modules [27]. From this analysis, we identified nine gene transcription modules and eight corresponding epithelial subclusters (Epi0-Epi7) (Figure 2A, Supplementary Data 2.1). The omentum-derived epithelial cells clustered together into Epi5 and Epi6, and the remaining clusters were composed of ovary-derived epithelial cells (Figure 2B). Each of the subclusters were transcriptionally different, as illustrated by the top 10 DEGs of each subcluster (Figure 2C, Supplementary Data 2.2). Similarly, GO enrichment analysis of the top DEGs revealed different functions for each gene transcription module (Supplementary Data 2.3-2.4) and subcluster (Figure 2D, Supplementary Data 2.5). Epi0 and Epi1 were enriched for epithelial morphogenesis terms; Epi2 and Epi4 were enriched for ECM and immune signaling terms; Epi3 was enriched for proliferation terms; Epi5 was enriched for differentiation, ECM, and proinflammatory signals terms; Epi6 was enriched for ECM and mesenchymal signaling terms; and Epi7 was enriched for cilium and axoneme terms. The module representation and cluster functions are summarized in Table 1.

**Figure 2:**
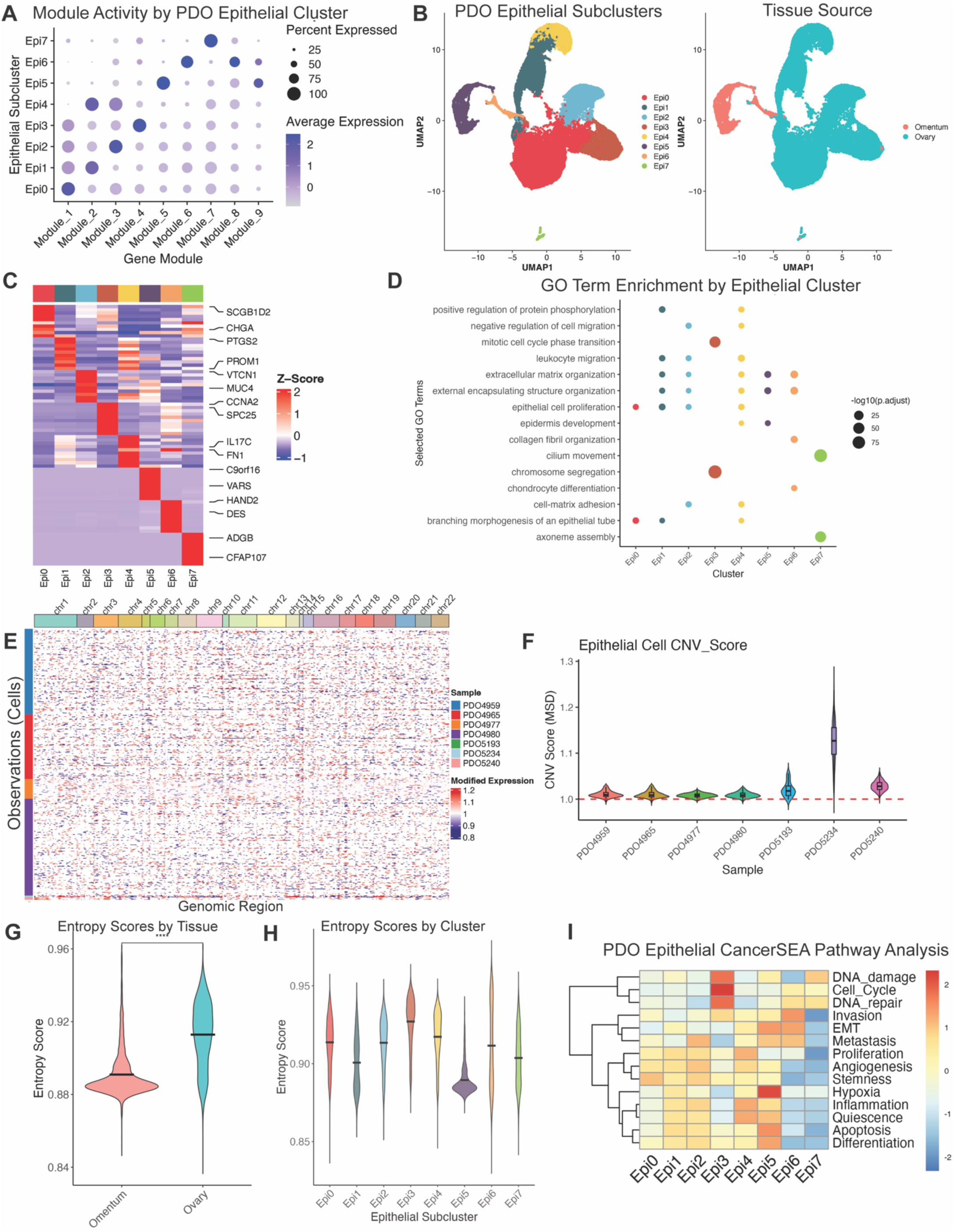
Epithelial cells from patient-derived organoids exhibit distinct transcriptional programs and functional states. A) Dot plot showing nine gene modules present in eight patient-derived organoid (PDO) epithelial subclusters (Epi0–Epi7). B) Uniform manifold approximation and projection (UMAP) visualization of epithelial cells, colored by subcluster identity and tissue source. C) Heatmap of top 10 marker genes defining each epithelial subcluster. D) Dot plot showing the top two Gene Ontology (GO) enrichment terms for differentially expressed genes in each epithelial subcluster. E) Heatmap displaying inferred large-scale copy number variation (CNV) profiles per epithelial cell. F) Violin plot of average CNV scores (mean squared deviation) across samples; outliers were excluded if they fell outside 1.5×interquartile range from the 1st and 3rd quartiles. CNV score >1 indicates high copy number variation. (G-H) Violin plots showing entropy scores stratified by (G) tissue source and (H) epithelial subcluster. I) Heatmap of PDO epithelial subclusters displaying tumor cell function gene set scores.

**Table 1.** Summary of PDO Epithelial Subcluster Functions.

| Cluster | Representative Modules | Cluster Function |
| --- | --- | --- |
| Epi0 | Module 1 | Epithelial Morphogenesis, Stress Response |
| Epi1 | Module 1, Module 2 | Progenitor, Epithelial Morphogenesis |
| Epi2 | Module 3, Module 1 | ECM, Immune-Modulatory |
| Epi3 | Module 4, Module 1 | Proliferative, Mitotic |
| Epi4 | Module 2, Module 3 | ECM, Inflammatory, Chemotactic |
| Epi5 | Module 5, Module 9 | Differentiation, Proinflammatory, ECM |
| Epi6 | Module 6, Module 8 | ECM, Mesenchymal-Primed |
| Epi7 | Module 7, Module 1 | Ciliated, Motile |

To assess whether PDO generation methods did not preferentially culture non-cancerous cells, we performed *inferCNV* analysis of copy number variation (CNV) in PDO epithelial cells. Genomic gains and losses were common across all chromosomes in each PDO sample (Figure 2E). The majority of PDO epithelial cells had a CNV score greater than 1, suggesting that most PDO epithelial cells were cancerous (Figure 2F). CNV scores of matched FFPE samples were similarly greater than 1 on average (Supplementary Figure 4).

We next performed single-cell entropy (SCENT) analysis to examine the cell differentiation potential and heterogeneity of PDO epithelial cells [28-30]. This analysis revealed that epithelial cells derived from ovarian tumors had significantly higher entropy scores than those derived from omental metastases (Figure 2G). Furthermore, the entropy scores of the epithelial cell subclusters significantly differed (Figure 2H). Epi 3, which was enriched for GO proliferative terms (Figure 2D), displayed the highest entropy score.

To further asses the functional differences across epithelial cell clusters, we performed gene set variation analysis (GSVA) using the CancerSEA database [31]. Each subcluster was enriched for distinct cancer-related pathways. For example, Epi3 showed the strongest association with DNA damage, cell cycle, and DNA repair pathways (Figure 2I). Taken together, these findings indicate that HGSOC PDO epithelial cells are functionally heterogeneous, with proliferative epithelial populations contributing disproportionately to higher entropy states.

### The PDO epithelial metastatic trajectory progresses from the ovary to the omentum

We next applied trajectory analyses common in stem cell and developmental biology analyses of non-cancerous organoids to examine changes associated with metastasis. First, we applied Potential of Heat-diffusion for Affinity-based Transition Embedding (PHATE) [32] to the PDO epithelial subcluster data to visualize cellular transitions in a two-dimensional space analogous to UMAP (Figure 3A). To model the transitions, we performed trajectory inference with Monocle3 [33] to reconstruct the continuum of transcriptional states associated with metastatic progression from primary ovary tumors to omental metastases. To define the root of the trajectory, we selected cluster Epi3, which came from ovary tumor samples (Figure 2B), displayed the highest entropy score (Figure 2H), and was the mitotically active (Supplementary Data 2.5). Initiating pseudotime from this cluster revealed a metastatic trajectory beginning with mitotically active ovary-derived epithelial cells (Ep3, lowest pseudotime) and terminating with omentum-derived cells (Epi5 and Epi6, highest pseudotime) (Figure 3B-C).

**Figure 3:**
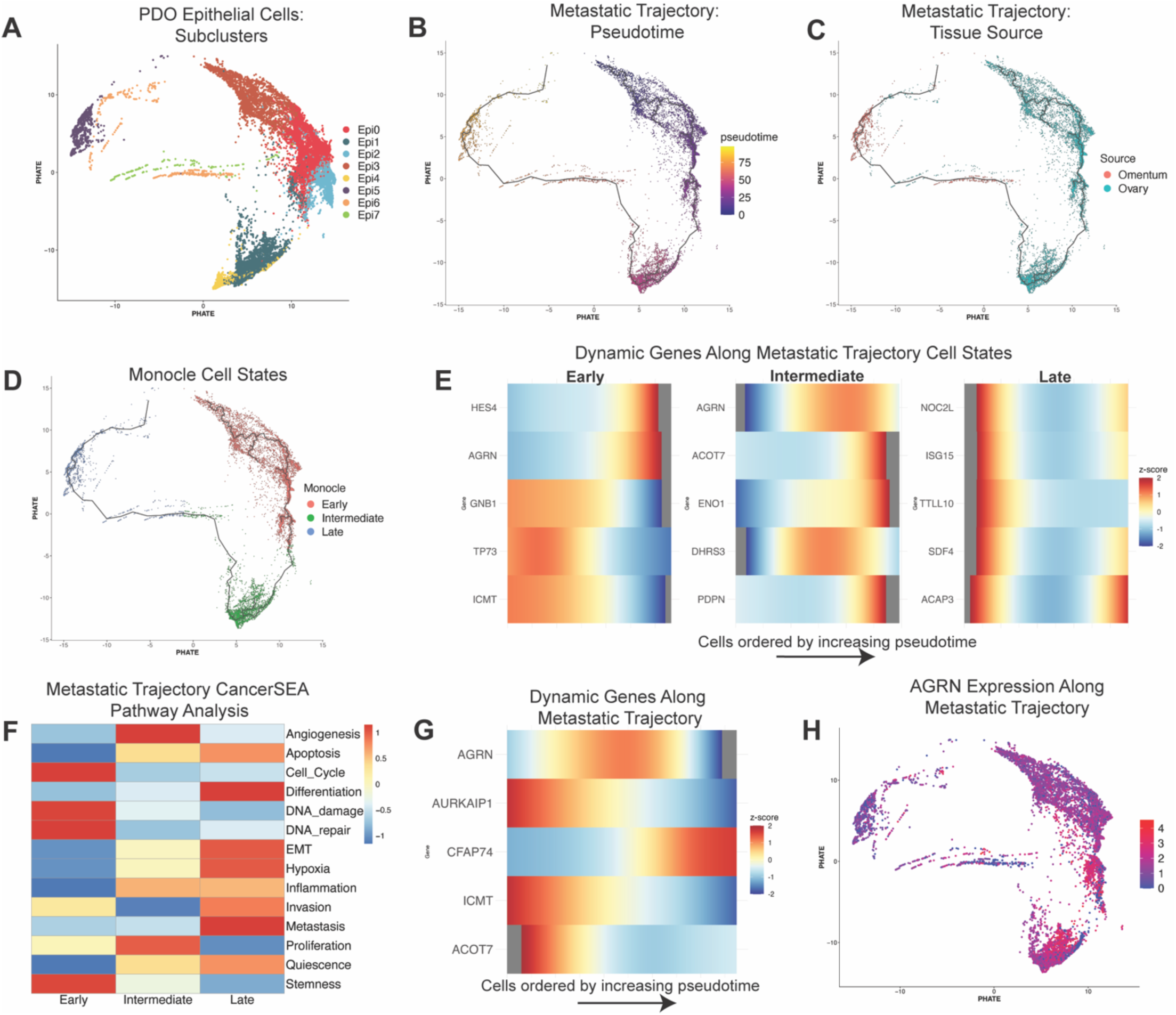
Pseudotime trajectory analysis of patient-derived organoid epithelial cells reveals distinct cell fate pathways and a metastatic progression axis. A) Potential of Heat-diffusion for Affinity-based Transition Embedding (PHATE) embedding of PDO epithelial cells colored by subclusters (Epi0–Epi7). B, C) Metastatic trajectory defined by pseudotime progression from ovary-derived to omentum-derived epithelial cells. PHATE plots depict epithelial cells annotated by (B) pseudotime and (C) tissue source. D) K-means clustering of metastatic trajectory of cells along the pseudotime axis stratified them into three distinct groups: early, intermediate, and late. E) Heatmaps of the top five differentially expressed genes per group. F) Gene Set Variation Analysis heatmap illustrating enrichment of CancerSEA pathways in early, intermediate, and late groups. G) Top five genes with significant spatial autocorrelation along the metastatic trajectory identified with Monocle3’s graph-based Moran’s I analysis. H) Metastatic trajectory feature plot of *AGRIN* expression.

Second, we applied k-means clustering to stratify cells into three monocle-defined transcriptional states: Early, Intermediate, and Late (Figure 3D). Using Monocle3’s graph-based Moran’s I analysis of each state, we identified the top 5 dynamically expressed genes associated with each state (Figure 3E). Each state was associated with different genes and expression patterns, which suggest state-specific transcriptional programs (Supplementary Data 3.1). Third, we performed GSVA CancerSEA analysis to assess whether the metastatic states were associated with specific cancer-related biological processes. The Early cells were most associated with stemness, DNA repair, DNA-damage, and cell cycle; the Intermediate cells were most associated with angiogenesis and proliferation; and the Late cells were most associated with invasion, metastasis and EMT (Figure 3F).

Fourth, to identify genes that may drive the metastatic trajectory, we used Monocle3 Graph_test analysis. This identified the top five genes whose expression changed the most over pseudotime (Figure 3G, Supplementary Data 3.2). One of these genes, which peaked in the Intermediate state (Figure 3H), was the heparan sulfate proteoglycan *AGRIN*. Although best known for its role in neuromuscular junction development [34], *AGRIN* promotes cellular proliferation, migration, oncogenic signaling, and tumor progression in hepatocellular carcinoma [35]. Additionally, it drives non–small cell lung cancer progression and enhances regulatory T-cell activation via IL-6 secretion through the PI3K/AKT pathway [36]. However, whether *AGRIN* contributes to HGSOC is unknown.

### PDOs exhibit greater *AGRIN*-driven intercellular signaling than matched FFPE samples

To identify intercellular signaling pathways that are more active or upregulated in PDOs than in matched FFPE tumor samples, we used CellChat [37]. PDO samples exhibited significantly more inferred interactions and stronger overall interactions than FFPE samples (Figure 4A, Supplementary Figure 5A). The outgoing and incoming signals were also higher in the PDO compared to FFPE samples (Supplementary Figure 5B). Pathway-level comparisons revealed enrichment of several metastasis-associated signaling routes in PDOs, including *Fibronectin 1* (FN1) [38] and *Collagen*. Consistent with our trajectory analyses (Figure 3G), *AGRIN* was the third highest pathway associated with metastasis (Figure 4B). *AGRIN* and most of the other top pathways showed higher signaling strength in PDOs than in FFPE tissues (Figure 4B). However, metastatic pathways such as *PECAM1* and *CDH5* [39] were higher in FFPE samples, suggesting that some signaling interactions are reduced or altered during the transition to *ex vivo* culture.

**Figure 4:**
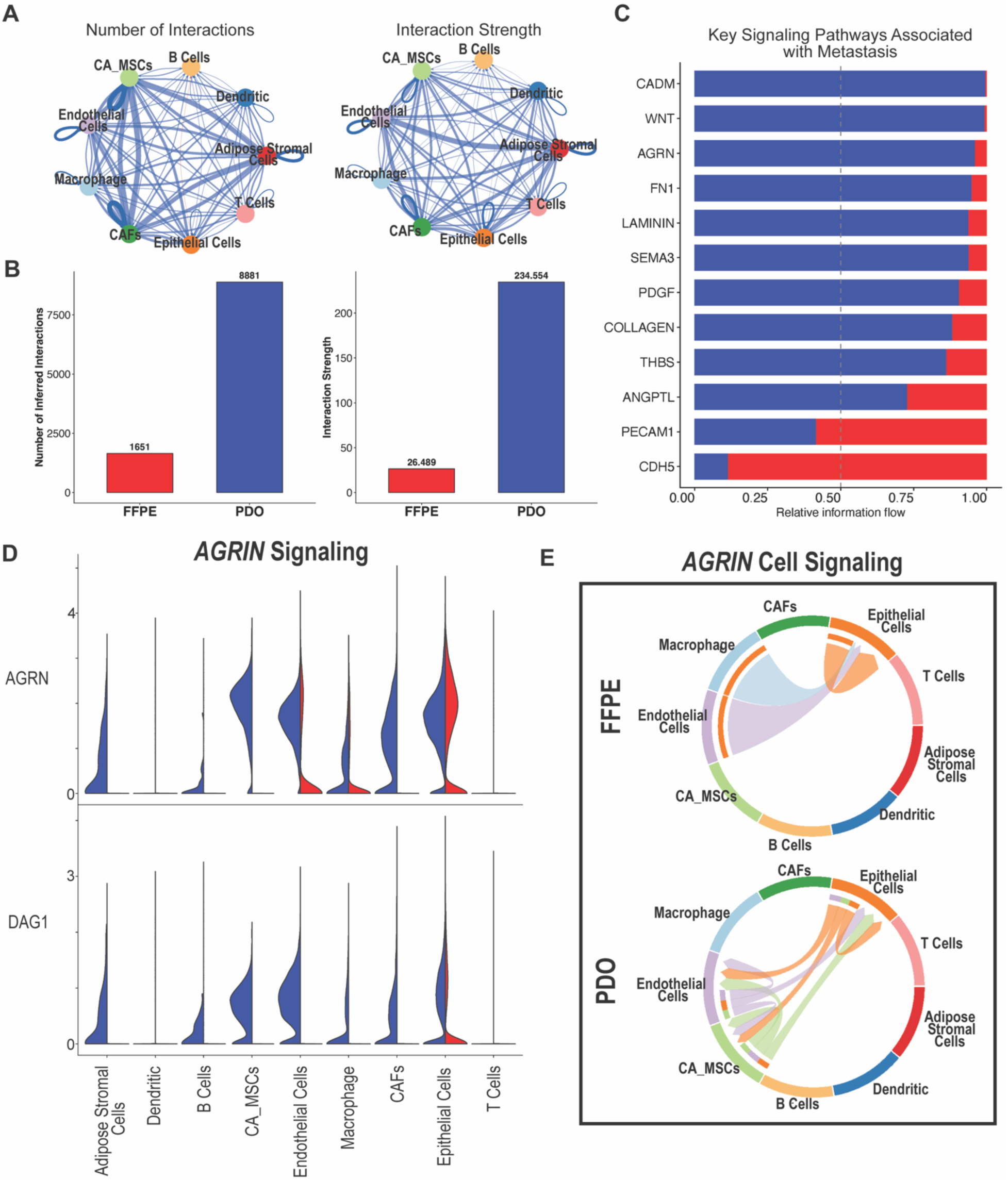
*AGRIN*-driven intercellular signaling is higher in patient-derived organoids than in matched frozen fixed paraffin-embedded tumor samples. A) Comparison of cell-cell communication networks between frozen fixed paraffin-embedded (FFPE, red) and patient-derived organoid (PDO, blue) samples. Circle plots and bar plot quantitation depicting numbers of interactions and interaction strengths. B) Relative information flow across 12 key metastasis-associated signaling pathways in PDO (blue) and FFPE (red) samples. C) Violin plots showing expression of *AGRIN* and *DAG1* in PDO (blue) and FFPE (red) samples. D) Chord plots illustrating *AGRIN* signaling network in PDO and FFPE samples. (AGRIN = AGRN)

Because *AGRIN* signaling emerged as a key feature in both our metastatic trajectory analysis and our cell–cell communication analysis, we focused further on this pathway. *AGRIN* signals through its receptor Dystroglycan (*DAG1*), a transmembrane glycoprotein that links the extracellular matrix to the cytoskeleton [40]. In PDOs, both *AGRIN* and *DAG1* were expressed in all cell types except dendritic cells and T cells (Figure 4C). In contrast, in FFPE samples, *AGRIN* was only expressed in endothelial cells, macrophages, and epithelial cells, whereas *DAG1* was largely restricted to epithelial cells. Chord diagrams depicting directional *AGRIN* signaling revealed that, in FFPE samples, *AGRIN*-mediated communication originated from macrophages, endothelial cells, and epithelial cells and was directed exclusively toward epithelial cells (Figure 4D). In PDOs, *AGRIN* signaling occurred to and from endothelial cells, epithelial cells, and CA_MSCs. Together, these findings suggest that *AGRIN* participates in multicellular signaling interactions spanning stromal, vascular, and immune-associated compartments, consistent with a broader role in shaping metastatic niche organization rather than acting solely as an epithelial-intrinsic regulator.

### AGRIN depletion suppresses invasive behavior in human HGSOC cells

Dysregulation of mechanotransduction in cancer contributes to increased tumor stiffness, disease progression, and enhanced metastatic potential [35, 41, 42]. AGRIN, an extracellular matrix heparan sulfate proteoglycan, strengthens integrin and DAG1 adhesion and promotes cytoskeletal tension, cell migration, and invasion [40, 43, 44]. Given these roles, we wondered whether AGRIN depletion would reduce metastatic behaviors in human ovarian cancer cell lines. To identify an appropriate cell line, we assessed AGRIN expression in OVCAR3, OVCAR8, and Kuramochi cells. OVCAR3 and Kuramochi exhibited detectable AGRIN expression, whereas OVCAR8 expressed minimal AGRIN (Figure 5A).

**Figure 5:**
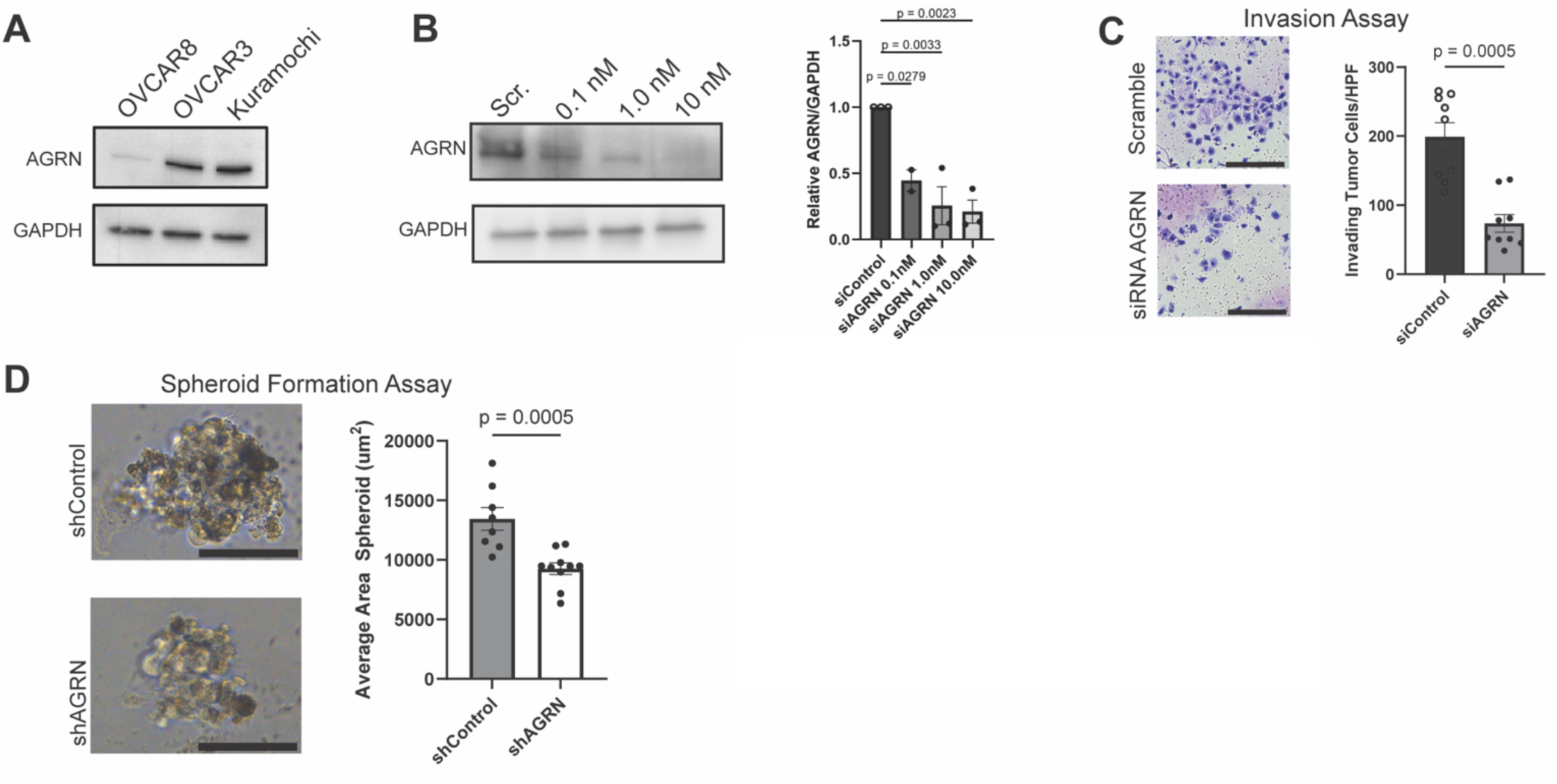
AGRIN depletion suppresses invasive and mesenchymal phenotypes in human high-grade serous ovarian cancer cells. A) Western blot analysis of AGRIN protein in human high-grade serous ovarian cancer epithelial cell lines OVCAR8, OVCAR3, and Kuramochi. GAPDH, loading control. B) Western blot and quantitation of AGRIN protein in Kuromochi cells 48 hours after transfection with *AGRIN* siRNA or scrambled control. GAPDH, loading control. C) Transwell invasion assay comparing scrambled and si*AGRIN*-transfected Kuramochi cells. Representative images and quantitation of stained invading cells 24 hours after seeding. D) Representative images and quantitation of spheroid formation in Kuramochi cells treated with *AGRIN* or Control shRNA. AGRIN = AGRN

RNAi-mediated (siRNA) depletion of *AGRIN* in Kuramochi resulted in greater than 50% reduction in protein expression (Figure 5B). To determine whether *AGRIN* loss impaired ovarian cancer cell invasion, we performed a transwell invasion assay in which we measured cell migration from serum-free media toward serum-rich media. In this assay, *AGRIN* depletion significantly reduced Kuramochi cell invasion (Figure 5C). Next, we assessed spheroid formation, which models anchorage-independent growth and multicellular aggregation, key features of metastatic dissemination in HGSOC. Kuramochi cells in which *AGRIN* was depleted formed smaller spheroids than shControl cells (Figure 5D).

## Discussion

Non-cancerous organoids, and the less heterogeneous spheroid models, have long been used in stem cell and developmental biology to study differentiation and niche dynamics. Here, we used a similar conceptual framework and analytical approaches to analyze HGSOC PDOs. Doing so allowed us to interrogate the complex, multicellular interactions that drive metastasis and to identify potential therapeutic targets. First, we performed scRNA-seq on matched PDOs and FFPE tumors. Within the first five passages, many epithelial, immune, and stromal cell types present in the matched FFPE tumors were retained in PDO cultures. However, we also observed loss of specific stromal subtypes, a reduction in immune populations, and a marked expansion of epithelial cells. Among the major cell compartments, PDO epithelial cells exhibited the greatest number of differentially expressed genes relative to matched FFPE samples. Despite these transcriptional changes, epithelial cells retained tumor-associated CNV patterns, indicating preservation of malignant identity. The relative depletion of immune populations in PDO cultures further suggests that these models preferentially capture tumor-intrinsic transcriptional programs and may underrepresent immune-dependent regulatory constraints that influence metastatic progression *in vivo*.

Second, entropy analysis revealed significant differences in cellular heterogeneity between ovarian- and omental-derived PDO epithelial cells. Trajectory analysis further suggested a continuum from ovary-derived to omentum-derived epithelial states, consistent with a metastatic progression axis. Because the omentum represents an immune-rich metastatic environment characterized by abundant macrophages, adipocytes, and stromal signaling networks [45-49], this transition likely reflects adaptation of tumor cells to a specialized secondary niche. We identified *AGRIN* as a candidate regulator of this transition. Cell–cell communication analysis supported this finding, demonstrating higher *AGRIN* signaling in PDOs than in matched FFPE samples. In addition, *AGRIN* inhibition in an HGSOC cell line reduced metastatic phenotypes. Although *AGRIN* has been implicated in metastasis in other malignancies, its role in HGSOC has not previously been established. Beyond its established functions in mechanotransduction and integrin-associated signaling, *AGRIN* has also been linked to cytokine-dependent pathways, including IL-6–mediated immune activation in other tumor contexts, suggesting that it may function at the interface of extracellular matrix remodeling and immune regulation within metastatic niches. Together, our data support *AGRIN* as a potential regulator of metastatic competence and a candidate therapeutic target in HGSOC.

Our work highlights that PDOs should not be viewed as miniature replicas of the original tumor but as evolving systems shaped by *in vitro* selective pressures. The expansion of epithelial populations and loss of certain stromal and immune compartments likely reflects differential survival and growth advantages under culture conditions. As our PDO culture medium did not contain exogenous cytokines, the immune cells in omentum-derived PDOs may be supported by cytokines and growth factors secreted by the TME. As PDOs are passaged, tumor-intrinsic programs that drive progression and metastasis may be selected for. Previous studies have demonstrated that PDOs recapitulate key genomic features and therapeutic drug sensitivities [50-56]. However, these models have primarily been utilized for testing candidate therapeutics rather than for systematic drug discovery.

A key strength of our study was the use of single-cell transcriptomics to uncover gene expression dynamics, as bulk analyses would have obscured shifts in subpopulation structures and cellular states. Additionally, entropy and trajectory analyses provided orthogonal evidence that PDO epithelial cells recapitulate biologically meaningful differentiation associated with metastatic progression. An important limitation of this study is the relatively small cohort size, consisting of seven matched PDO–FFPE pairs derived from patients previously treated with chemotherapy. As a result, the observed tumor microenvironmental changes may reflect, in part, therapy-associated remodeling. Future studies should validate these findings in larger, treatment-diverse cohorts, including chemotherapy-naïve samples, to better define the intrinsic and treatment-associated features of HGSOC progression.

In summary, this study evaluates the utility of HGSOC PDOs as a platform for identifying novel molecular and therapeutic targets. Despite limitations, including a modest sample size of seven matched PDO–FFPE pairs, our results support the concept that living tissue biobanks coupled with high-resolution transcriptomics can generate functional “digital twins” of patient tumors. Importantly, discrepancies between PDOs and primary tissues, rather than being viewed solely as artifacts, can serve as a source of discovery. Our findings provide a framework for using PDOs as dynamic models to uncover regulators of tumor progression and metastasis, supporting their application in future molecular studies and therapeutic development. More broadly, our findings support a model in which AGRIN functions within interconnected extracellular matrix, stromal, and immune signaling networks that collectively shape metastatic competence, highlighting the value of single-cell approaches for identifying regulators of tumor progression within complex multicellular microenvironments.

## Methods

### Patient-derived Organoid Generation

All human tissue and specimens were collected in accordance with Washington University in St. Louis Institutional Review Board-approved protocols. PDOs were generated as described [57]. In short, primary and metastatic ovarian tumor tissue was obtained at the time of debulking surgery. Samples were mechanically and enzymatically dissociated in Base Media (Advanced DMEM/F12, 1× Glutamax, 1 mg/mL Primocin, 1% HEPES) containing 1 mg/mL type III collagenase and 100 µg/mL DNase for 1 hour at 37 °C in a gentleMACS Tissue Dissociator. The dissociated tumors were filtered through a 100 µm strainer and centrifuged at 1,650 x g for 5 min at 4 °C. Cells were then resuspended in Cultrex Type 2 and treated with M2 media, supplemented with growth factors as described [8]. PDOs were passaged every 7–10 days with TrypLE Express and maintained at 37 °C with 5% CO₂. For scRNA-seq, early-passage organoids (P1–P7) were used.

### FFPE Generation

Bulk tumor tissue pieces matched to PDOs were fixed in 10% neutral-buffered formalin for 24 hours at room temperature, processed through graded ethanol dehydration steps, and embedded in paraffin. Tissues blocks were stored at room temperature until sectioning.

### scRNA Sequencing Sample Preparation

For FFPE samples, four 25 µm curls (per sample) were cut with a microtome, deparaffinized with xylene, and sequentially rehydrated in 100%, 95%, and 70% ethanol. Samples were then processed according to a standardized FFPE tissue dissociation workflow (10x Genomics, CG000632) yielding single-cell suspensions. For PDOs, cells were disassociated, fixed, and processed according to a tissue dissociation workflow (10x Genomics, CG000553). FFPE- and PDO-derived single-cell suspensions were resuspended in PBS containing 0.04% BSA and filtered through a 40 µm strainer. AO/PI staining and fluorescent cell counting (Revvity K2) were used to determine cell number. were used for library generation. Samples were diluted to a target concentration of 700–1,200 cells/µL and loaded into the Chromium Controller (10x Genomics) by using the Chromium Single Cell 3’ v3.1 or 5’ v1.1 chemistry according to the manufacturer’s specifications. The target cell recovery per sample was 8,000–10,000 cells.

### scRNA Sequencing

Gel bead-in-emulsion generation, cell lysis, reverse transcription, and cDNA amplification were carried out according to 10x Genomics guidelines. Amplified cDNA was quantified with a Qubit fluorometer and assessed with an Agilent Bioanalyzer. Sequencing libraries were prepared via enzymatic fragmentation, end repair, A-tailing, adapter ligation, and sample indexing. Libraries were pooled at equimolar ratios and sequenced on an Illumina NovaSeq 6000 platform using S4 flow cells to achieve an average depth of 40,000–50,000 reads per cell. Library structure included a 16 bp cell barcode, 10 bp unique molecular identifiers, and 28 bp Read 1 sequence. FASTQ files were processed with CellRanger (v7.x) using the GRCh38 reference transcriptome. Feature-barcode matrices were used for downstream analysis.

### Sample Quality Control and Integration

Raw matrices were loaded into Seurat (v4.3.0). Cells were filtered to include those with >500 detected genes, >1,000 unique molecular identifiers, and <20–30% mitochondrial content. Doublets were removed by using DoubletFinder with an expected doublet rate of 5–7% based on the number of recovered cells. Data were normalized with SCTransform, and variable features were selected by using the VST method (2,000 genes). Samples were integrated with Harmony to correct for batch- and sample-level variation. Principal component analysis was performed on scaled data, and uniform manifold approximation and projection embeddings were computed from the top 30 principal components.

### Differential Gene Expression Analysis

Cell types were pseduobulked into four groups: epithelial, endothelial, stromal, and immune cells. Differential gene expression between cell groups and sample type (PDO vs. FFPE) was computed by using the DESeq2 function with the Wilcoxon rank-sum test. Only genes expressed in >10% of cells per group and with adjusted p-value <0.05 (Bonferroni) were considered significant. Top marker genes were used to annotate clusters and define cell groups.

### Non-negative Matrix Factorization (NMF) Analysis

NMF was performed to identify gene co-expression modules across epithelial subsets. The log-normalized matrix was decomposed in the NMF R package (v0.23.0), testing ranks k = 4–12. Optimal k was selected by using cophenetic correlation and dispersion metrics. Module scores were calculated per cell and visualized across pseudotime and epithelial states.

### Monocle3 Analysis

Trajectory inference was performed with Monocle3 (v1.2.9). The integrated Seurat object was converted to a cell_data_set object by using SeuratWrappers. The learn_graph function was used to generate the principal graph. The root node for pseudotime calculation was assigned to the epithelial subcluster with the highest entropy score and expression of stem-like markers (Epi3). Pseudotime values were extracted and used for downstream visualization and gene trend analysis.

### Single-Cell Entropy (SCENT) Analysis

Cellular entropy scores were computed by using the SCENT algorithm to estimate transcriptional plasticity across epithelial subsets. Raw expression matrices were normalized as recommended by the SCENT pipeline. Entropy per cell was calculated based on signaling network flux estimations mapped to the human protein interaction network. Subclusters with elevated entropy were operationally defined as highly plastic or stem-like populations.

### Copy Number Variation (CNV) Analysis

To assess malignant transformation in epithelial clusters, CNV profiles were inferred by using inferCNV (v1.12). Normal fibroblasts and endothelial cells served as reference “normal” populations. Raw counts were used as input, and genes were ordered according to their genomic coordinates. The denoised CNV signal was generated by using the default Hidden Markov Model and Bayesian smoothing. Clusters exhibiting broad copy number gains or losses characteristic of HGSOC (e.g., gains in chromosomes 3q, 8q, or 20q; losses in chromosomes 4, 5q, or 16q) were classified as malignant epithelial cells.

### Gene Set Variation Analysis (GSVA)

The GSVA package (v1.42) was used with CancerSEA functional signatures, including stemness, epithelial-to-mesenchymal transition, metastasis, angiogenesis, DNA repair, and cell cycle. GSVA enrichment scores were compared across pseudotime-defined cell states (Early, Intermediate, Late) by using Wilcoxon rank-sum tests. Pathways with false discovery rate <0.05 were considered significant.

### Cell-Cell Interaction Analysis

Cell–cell communication networks were inferred with CellChat (v1.6.1). Normalized expression matrices from integrated PDO and FFPE datasets were analyzed separately. Significant ligand–receptor interactions were identified by using the database-curated signaling lists.

Communication probability was estimated via permutation testing with 1,000 iterations. Overall interaction strength, number of interactions, and pathway-specific contributions were compared between PDO and FFPE samples. Selected pathways were visualized by using netVisual_circle and chord diagrams.

### Cell Lines

The human ovarian cancer epithelial cell line Kuramochi were obtained from ATCC. Cells were maintained in RPMI-1640 supplemented with 10% fetal bovine serum, 1% penicillin–streptomycin, and 2 mM L-glutamine. Pooled human umbilical vein endothelial cells (HUVECs; Lonza) were cultured in EGM-2 BulletKit medium (Lonza). All cell lines were maintained at 37 °C in 5% CO₂ and routinely tested for Mycoplasma contamination.

### Migration Assay

Transwell migration was assessed with uncoated inserts. In short, 50,000 serum-starved cells were seeded into the upper chamber in serum-free medium and allowed to migrate toward media supplemented with 10% FBS for 24 hours. Migrated cells were fixed, stained (Hema 3 Stat Pack, Fisherbrand), and quantified by using the ImageJ Cell Count plugin. For AGRIN knockdown conditions, siRNA (IDT) was transfected 24 hours before assay initiation.

### Conditioned Media Experiments

HUVECs at passages 2–5 were grown to confluence and then incubated for 72 hours in either complete EGM-2 medium (containing all BulletKit supplements) or basal EGM-2 medium supplemented with FBS only (lacking growth factor supplements). Conditioned media was collected and centrifuged at 4,000 × g for 10 minutes to remove debris. EGM-2 conditioned media or corresponding non-conditioned control media was mixed with epithelial cell culture media at a 40:60 ratio. Epithelial cells were incubated under these conditions for 48 hours before protein isolation.

### Spheroid Formation

Two hundred Kuramochi cells were seeded into low-attachment 96-well plates in quintuplicate. After 7 days in Kuramochi cell media, spheroid formation was assessed by microscopy, and average spheroid area was quantified in ImageJ.

### Western Blotting

Cells were lysed in RIPA buffer supplemented with protease inhibitors. Protein concentration was quantified with the BCA assay. Equal amounts of protein (15-20 µg) were separated via SDS–PAGE (10%) and transferred to PVDF membranes. Membranes were blocked in 5% milk and incubated overnight at 4 °C with primary antibodies: anti-AGRIN (ProteinTech; 1:500), anti-GAPDH (Cell Signaling; 1:1,000), and anti-Vimentin (Cell Signaling, 1:1000). HRP-conjugated secondary antibodies were applied for 1 h at room temperature. Signal was detected with Enhanced Chemiluminescent reagent (MilliporeSigma) and imaged on a ChemiDoc MP system.

