## Supplemental Figures1-5 for "Single-cell transcriptomics of heterogeneous patient-derived organoids reveals novel therapeutic targets in high-grade serous ovarian cancer"

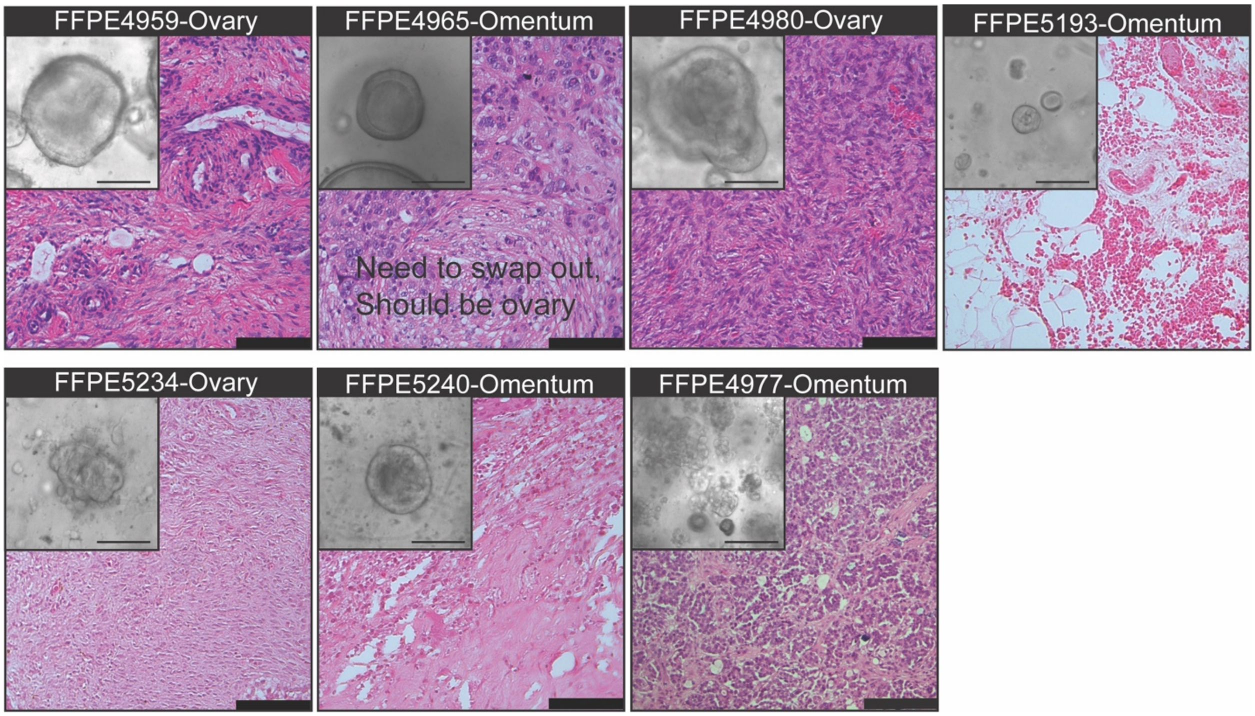


**Supplementary Figure 1:** A) Brightfield hematoxylin and eosin (H&E) images show matched FFPE tumor sections at 20× magnification with a scale bar of 100 μm. Insets show corresponding PDOs at the time of sequencing imaged by brightfield microscopy at 20× magnification with a scale bar of 25 μm.

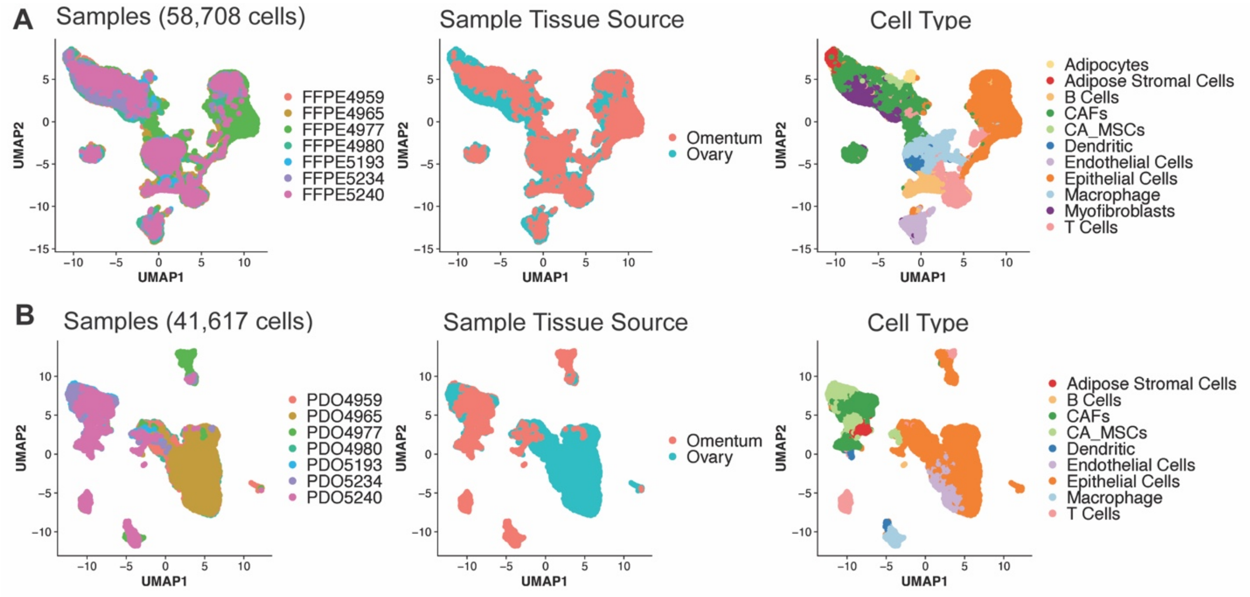


**Supplementary Figure 2:** A) UMAP embeddings of FFPE samples colored by individual sample identity, sample tissue source, and annotated cell type. B) UMAP embeddings of PDO samples colored by individual sample identity, sample tissue source, and annotated cell type.

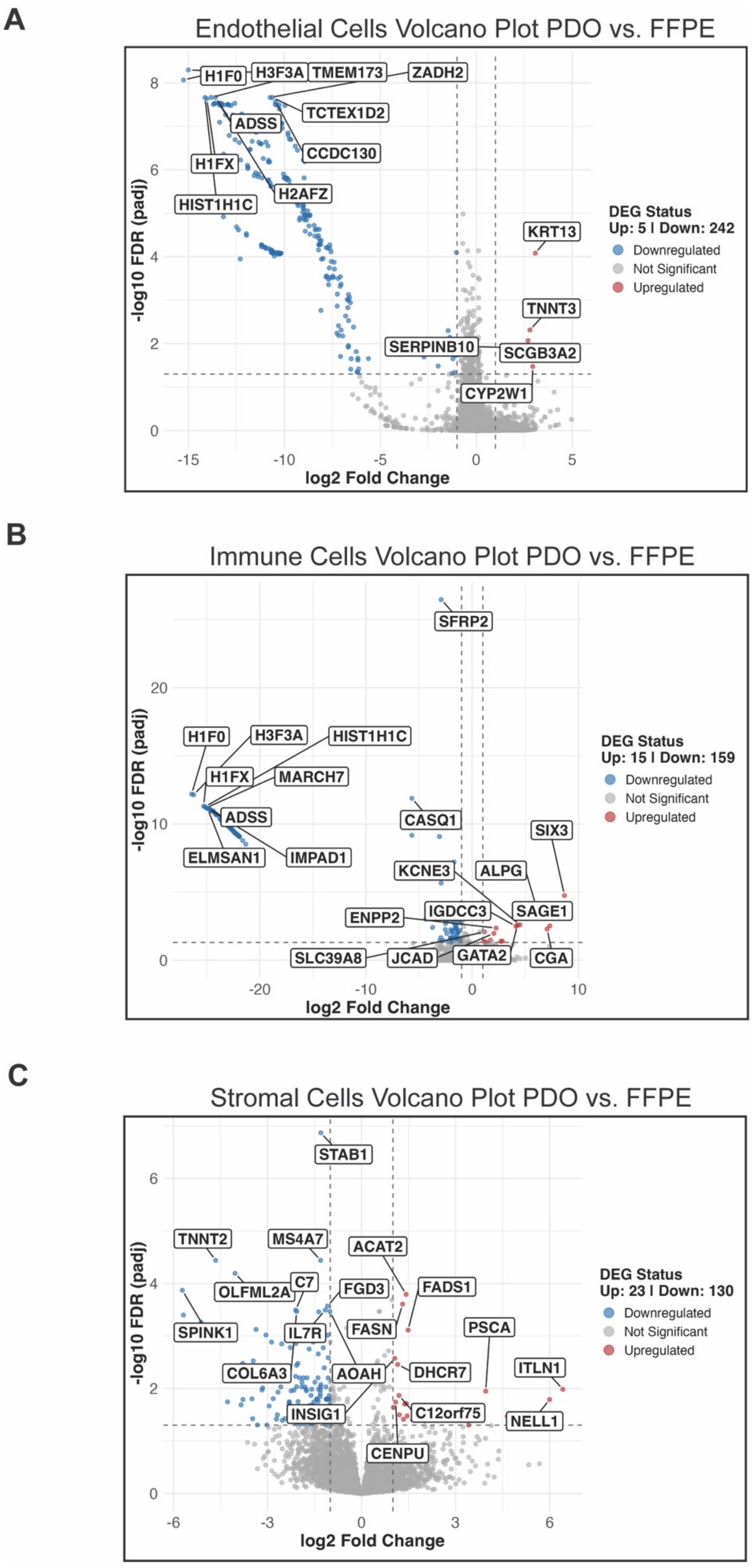
A
Endothelial Cells Volcano Plot PDO vs. FFPE
-log10 FDR (padj)
log2 Fold Change
H1F0
H3F3A
TMEM173
ZADH2
TCTEX1D2
ADSS
CCDC130
H1FX
H2AFZ
HIST1H1C
KRT13
TNNT3
SERPINB10
SCGB3A2
CYP2W1
DEG Status
Up: 5 | Down: 242
Downregulated
Not Significant
Upregulated
B
Immune Cells Volcano Plot PDO vs. FFPE
-log10 FDR (padj)
log2 Fold Change
SFRP2
H1F0
H3F3A
HIST1H1C
H1FX
MARCH7
ADSS
ELMSAN1
IMPAD1
CASQ1
SIX3
KCNE3
ALPG
IGDCC3
SAGE1
ENPP2
SLC39A8
JCAD
GATA2
CGA
DEG Status
Up: 15 | Down: 159
Downregulated
Not Significant
Upregulated
C
Stromal Cells Volcano Plot PDO vs. FFPE
-log10 FDR (padj)
log2 Fold Change
STAB1
TNNT2
MS4A7
ACAT2
C7
OLFML2A
FGD3
FADS1
SPINK1
IL7R
FASN
PSCA
COL6A3
AOAH
DHCR7
ITLN1
INSIG1
C12orf75
NELL1
CENPU
DEG Status
Up: 23 | Down: 130
Downregulated
Not Significant
Upregulated

**Supplementary Figure 3:** Volcano plots displaying differentially expressed genes (DEGs) between PDO and FFPE samples for A) endothelial cells, B) immune cells, and C) stromal cells. The x-axis represents log2 fold change (PDO vs. FFPE), and the y-axis represents –log10 adjusted p-value.

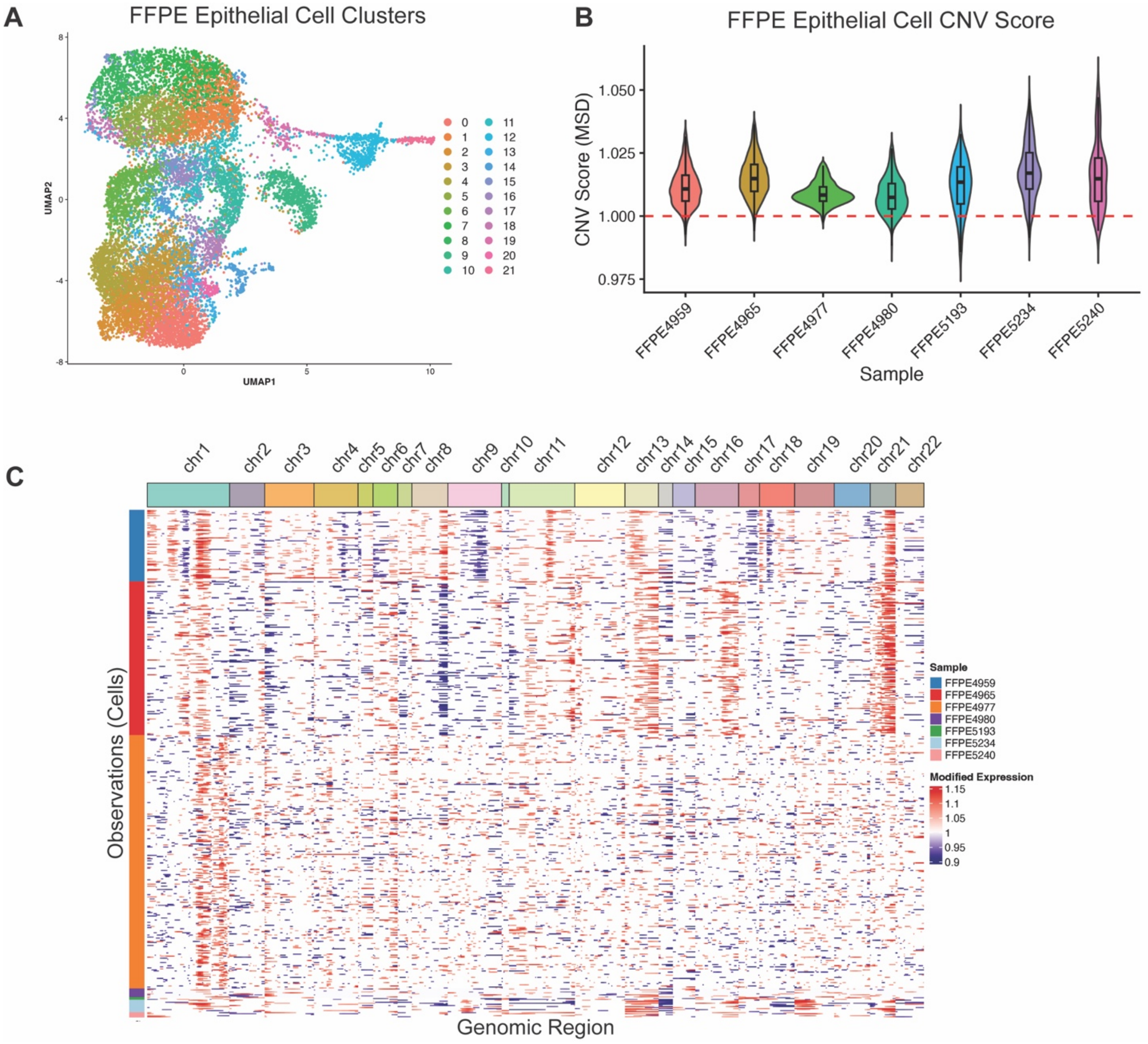


**Supplementary Figure 4:** A) The UMAP plot shows FFPE epithelial cells colored by cluster identity. B) The violin plot shows inferred CNV scores across FFPE samples, with the red line indicating a CNV score of 1.0 and values greater than 1 indicating increased copy number variation. C) The heatmap shows inferred large-scale copy number variation profiles across all epithelial cells.

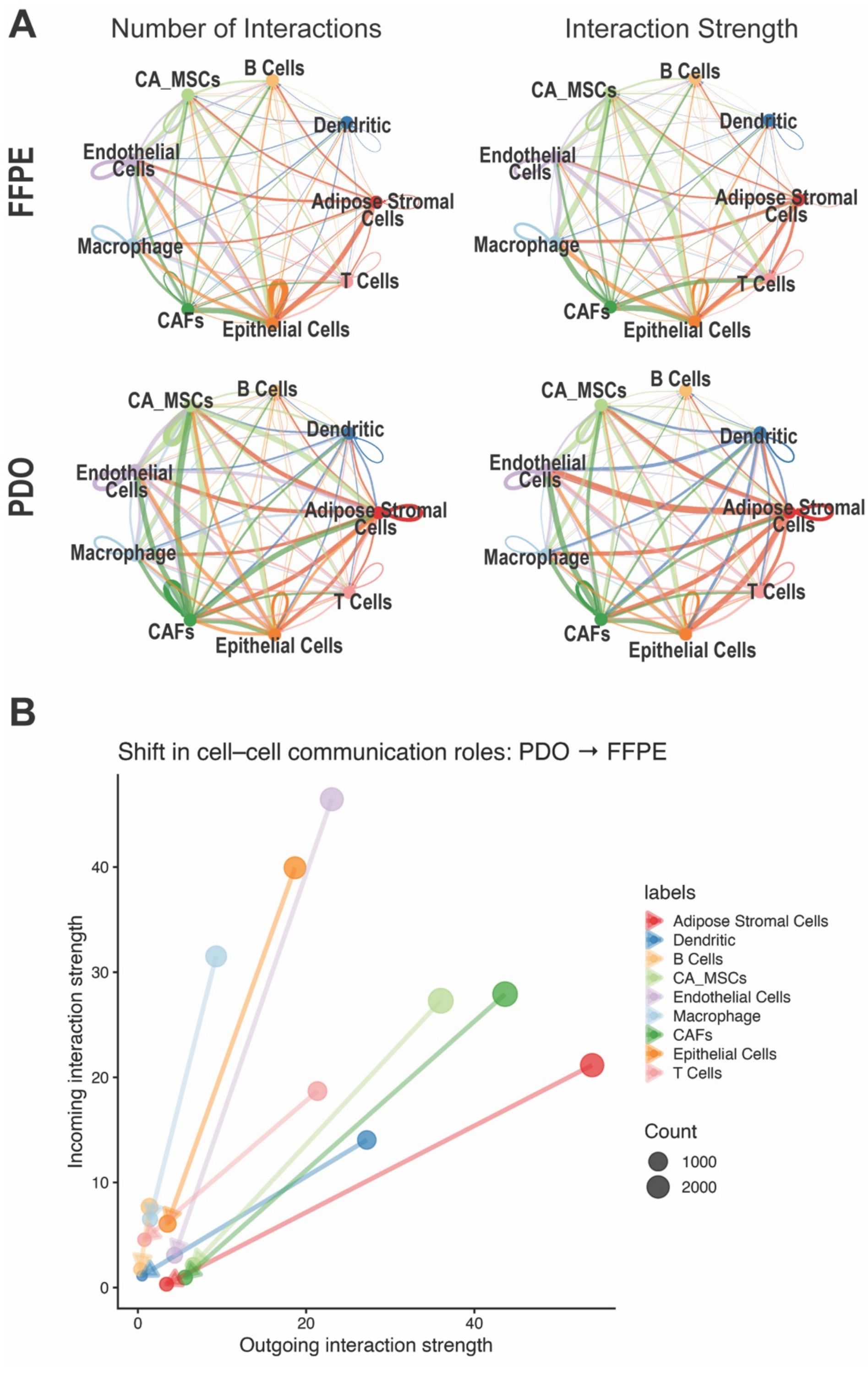


**Supplementary Figure 5:** A) The circle plot shows the number of interactions and interaction strength between cell types in FFPE and PDO samples as inferred by CellChat. B) The plot shows shifts in cell–cell communication roles from PDO to FFPE, with arrows indicating directional changes in signaling roles between conditions.
